# Intestinal parasites among rural school children in southern Ethiopia: A cross-sectional multilevel and zero-inflated regression model

**DOI:** 10.1101/2019.12.19.882217

**Authors:** Hiwot Hailu, Bernt Lindtjørn

**Affiliations:** School of Public Health, College of Medicine and Health Sciences, Hawassa University, Hawassa, Ethiopia; Centre for International Health, University of Bergen, Bergen, Norway; Department of Public Health, College of Health Sciences and Medicine, Dilla University, Dilla, Ethiopia

## Abstract

**Background:** Over 28 million school-aged children are at risk of intestinal parasite infection in Ethiopia. Few studies have investigated household-level risk factors or applied multilevel analysis to account for the nested data structure. This study aimed to assess the prevalence, intensity, and risk factors of parasite infection among schoolchildren in rural South Ethiopia.

**Methodology/Principal Findings:** Using multistage random sampling, we recruited 864 students in the Wonago district. We applied multilevel-logistic and zero-inflated negative binomial regression models (ZINB). Risk factors were concentrated at the individual level; school-level and class-level variables explained less than 5% of the variance. The overall intestinal parasite prevalence was 56% (479/850); *Trichuris trichiura* prevalence was 75.2% (360/479); and *Ascaris lumbricoides* prevalence was 33.2% (159/479). The rate of infection increased among children with anemia (AOR: 1.45 [95% CI: 1.04, 2.03]), wasting (AOR: 1.73 [95% CI: (1.04, 2.90]), mothers who had no formal education (AOR: 1.08 [95% CI: 1.25, 3.47]), and those in households using open containers for water storage (AOR: 2.06 [95% CI: 1.07, 3.99]). In the ZINB model, *A. lumbricoides* infection intensity increased with increasing age (AOR: 1.08 [95% CI: 1.01, 1.16]) and unclean fingernails (AOR: 1.47 [95% CI: 1.07, 2.03]). Handwashing with soap (AOR: 0.68 [95% CI: 0.48, 0.95]), de-worming treatment [AOR: 0.57 (95% CI: 0.33, 0.98)], and using water from protected sources [AOR: 0.46 (95% CI: 0.28, 0.77)] were found to be protective against parasitic infection.

**Conclusions/Significance:** After controlling for clustering effects at the school and class levels and accounting for excess zeros in fecal egg counts, we found an association between parasite infections and the following variables: age, wasting, anemia, unclean fingernails, handwashing, de-worming treatment, mother’s education, household water source, and water storage protection. Improving hygiene behavior, providing safe water at school and home, and strengthening de-worming programs is required to improve the health of schoolchildren in rural Gedeo.

**Author summary:** Intestinal parasite infections are common among school-aged children in Ethiopia. Several cross-sectional studies have investigated the prevalence and risk factors of these intestinal parasite infections. However, most were conducted in an urban setting in northern Ethiopia; they collected household-level risk factor information from the children, not the parents; and they restricted intestinal parasite infection data to binary outcomes. Therefore, we aimed to assess the prevalence and intensity of intestinal parasite infections and the related individual-, household-, and school-level risk factors among rural schoolchildren in southern Ethiopia. Using a multivariate, multilevel, regression model, we found minimal variation across class- and school-level factors for intestinal parasite infection prevalence. We found associations between intestinal parasite infections and most individual-level factors and some household-level factors. Therefore, interventions focusing on the individual, household, and school should be implemented to reduce the prevalence of infection and parasite load among schoolchildren.

## Introduction

More than 1.5 billion people around the world are affected by intestinal parasites, including over 568 million schoolchildren who are at risk (1). In 2015, an approximately 88 million individuals, including 28 million school-aged children, were at risk for intestinal parasite infection in Ethiopia (2). Roundworm (*Ascaris lumbricoides)*, whipworm (*Trichuris trichiura*), and hookworm (*Ancylostoma duodenale* and *Necator americanus*) are the most common intestinal helminth infections that chronically infect children (3). Neglected tropical diseases tend to receive less attention and funding than malaria or TB research (3, 4), although the economic and health burdens of these infections exceed those for other diseases (4, 5). Anemia, poor physical and intellectual development, and impaired cognitive function can occur with these infections (6, 7). The risk factors of intestinal parasite infections include poverty (8), mothers’ education, untrimmed fingernails, walking barefoot, unsanitary toilet areas, not washing hands before eating or after visiting the toilet, eating raw or undercooked vegetables or meat, lack of hygiene facilities, and drinking water from unsafe sources (9). In developing countries, control measures can be difficult to implement due to water and sanitation problems (10).

In Ethiopia, the prevalence of intestinal parasite infection among schoolchildren ranges between 18% and 81% (8, 11-17), with the highest rate of infection (81%) recorded in the southern region (17). The government of Ethiopia is expanding schooling to make education more relevant to all children and meet their nutritional and health needs (18). This strategy includes facilitating and implementing a de-worming service every six months and improving water, hygiene, and sanitation facilities (19). However, many school-aged children continue to be affected by parasitic infections (20-22). Moreover, most schools have no handwashing facilities, and hygienic behavior is inadequate (23).

School-based, cross-sectional studies have been used to study the prevalence and predictors of intestinal parasite infections (8, 11-17). Many of these studies have been conducted in urban settings in northern Ethiopia, and they focused on school feeding programs (24) and infection prevalence (25). Unfortunately, they did not assess potential household risk factors or adequately address the nested structure of school data (i.e., individuals nested within the same classes, which are nested within the same schools). Furthermore, most interpreted the intestinal parasite infection data in terms of binary outcomes (e.g., presence and absence).

Few studies have assessed the prevalence of intestinal parasite infections in southern Ethiopia (17). This paper therefore aimed to assess the prevalence and intensity of intestinal parasite infections among rural school children in the Wonago District of South Ethiopia, as well as the individual-, household-, and school-level factors that contribute to these infections. In addition to the binary outcome variable, we considered intestinal parasite egg concentration in the stool specimen. Using a multivariate, multilevel, regression model, we identified factors contributing to variations in the prevalence of intestinal parasite infections in this population.

## Method

### Study design and setting

We conducted this cross-sectional study from February 2017 to June 2017 in the Wonago district of the Gedeo Zone in the southern region of Ethiopia. The district is 375 km south of Addis Ababa, the capital city of Ethiopia. The district has 17 rural and 4 urban *kebeles*, which is the smallest administrative unit. In 2014, Wonago’s population was estimated at 143,989 people: 71,663 (49.8%) men and 72,326 (50.2%) women. The population density of the district is 1,014 persons per square kilometer, making it one of the most densely populated areas in Ethiopia. The district has 26 government health facilities (6 health centers and 20 health posts), 2 private clinics and 2 drug stores, and more than 36,000 students in 3 urban and 22 rural primary schools. Most residents depend on cash crops of coffee, fruits, and *ensete* (*Ensete ventricosum).*

### Participants

We recruited students aged 5–14 years who gave consent and whose parents or guardians gave consent to participate. Using a three-stage cluster sampling method, we assigned schools to level one, classes to level two, and students to level three. We replaced participants who dropped out of school after the selection process with participants of similar class, sex, and age. We selected only one child per household and collected household information from the parent or guardian. Figure 1 shows the recruitment process and participant profile.

**Figure 1:**
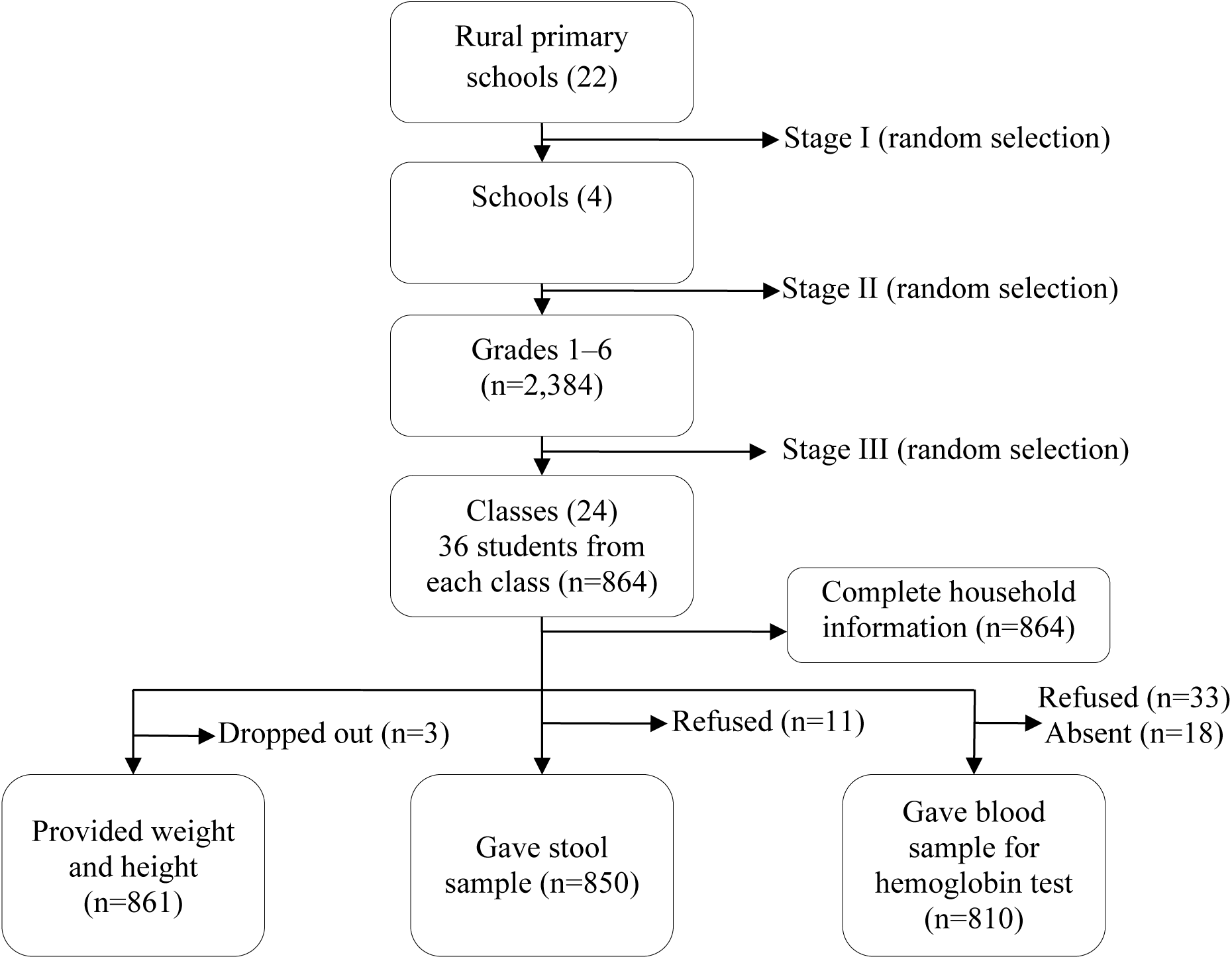
Study recruitment profile, Wonago district, Gedeo zone, southern Ethiopia, 2017.

### Sample size

We calculated the sample size using OpenEpi software (26), assuming a 95% confidence interval (CI), maximum sample sizes based on proportions of different variables (e.g., underweight [59.7%], stunting [30.7%], wasting [37.2%], intestinal parasite [27.7%], and skin infection [50%]), 5% precision, and design effect 2 due to multistage sampling. We also calculated the sample size using outcome-associated variables, such as undernutrition (32.2% prevalence of stunting among female participants), intestinal parasites (52.6% among children who did not wash their hands before meals), and skin infection (21.8% among children aged 6–10 years). To yield the maximum sample size, we calculated 50% for skin infection (12, 27-29). After adding a 10% non-response rate, we reached a final sample size of 845, the minimum required sample size. We then randomly recruited 864 students.

### Variables and measurements

We used a multilevel logistic regression model to analyze three separate binary outcome variables: the presence or absence of any intestinal parasite infections, the presence or absence of *T. trichiura*, and the presence or absence of *A. lumbricoides*. Two separate count models also were constructed for the *T. trichiura* and *A. lumbricoides* fecal egg counts. The association between the over-dispersed count outcome with excess zeros and the potential predictors of *T. trichiura* and *A. lumbricoides* infection was determined using a zero-inflated negative binomial (ZINB) regression model (30).

The intensity of parasite infection was measured according to density of eggs in stool samples. Using the Kato-Katz technique, each parasite egg was multiplied by 24 to quantify the result in eggs per gram (epg) of stool (31). Infection intensity was defined as light (<5,000 epg for *A. lumbricoides;* <1,000 epg for *T. trichiura;* and <2,000 epg for hookworm) or moderate (≥5,000 epg for *A. lumbricoides*; ≥1,000 epg for *T. trichiura*; and ≥2,000 epg for hookworm) (32).

We assessed individual, parent, and household exposure variables. The individual child factors included sex, age, hygiene behavior, loss of appetite in the past month, de-worming treatment in the past 6 months, anemia (< 11.5g/dl for those aged 5–11 and < 12g/dl for those aged 12–14), stunting (height-for-age Z scores below −2 SD), and wasting (body-mass-index-for-age Z scores below −2 SD). Parent factors included the educational level of the mother and father. Household factors included the wealth index, which was constructed using principal component analysis of 15 household assets (electricity, radio, television, mobile phone, table, chair, bed, separate kitchen, cooking place, own land, bank account, toilet facility, floor type, and roof type); family size; source of drinking water; container used to store water; and use of treated water. School factors included access to health education on personal hygiene, absence from school in the past month, and participation in the school food program.

Stool samples were collected, processed, and examined using standard procedures (33, 34). Samples were collected in the early morning at school and stored in stool cups labeled with an identification code, name, sex, age, and date. Specimens then were transported in a cold-box with frozen ice-packs to the nearest health facility, Dilla University Teaching and Referral Hospital, where Kato-Katz and formalin-ether concentration techniques were used to conduct stool tests. Single, 41.7-mg thick, Kato-Katz smears were prepared from each stool sample on the same day of specimen collection. Then, 1 g of stool was preserved in 10% formalin solution and processed using the formalin-ether concentration technique (35). The slides were examined by three experienced laboratory technicians. The result of each parasite species from two diagnostic techniques was recorded separately. Individual and household factors were collected from the child or the child’s parents or guardians via interviews and observations of housing conditions. Hemoglobin concentrations were measured from capillary blood samples using a HemoCue Analyser Hb 301 (Angelholm, Sweden) (36). Anthropometric indices for height-for-age and body-mass-index-for-age also were calculated.

### Data quality control and statistical methods

To minimize potential bias and validate the measurement tools prior to actual data collection, a pre-test was conducted on 42 primary schoolchildren in other schools not selected for this study. Ten trained enumerators used a pretested, structured questionnaire that was primarily adapted and developed in English and then translated into the local language (*Gedeooffa*). Supervisors checked the data onsite for completeness and consistency.

All Kato-Katz slides were examined within one hour of preparation to minimize bias. To reduce possible bias introduced during outcome measurement, 10% of the test results were re-examined in a blinded fashion to ensure reproducibility of the results. Kappa statistics (STATA 14 software) were used to estimate reliability of the inter-rater agreement of the two readers. Kappa values were defined as follows: poor=0.01–0.2; fair=0.21–0.4; moderate=0.41–0.6; good=0.61–0.8; and perfect=0.81–1 (37). Kappa values were considered statistically at P <0.05. The reliability (Kappa values) were as follows: *A. lumbricoides*, 0.83 [0.72 – 0.95; 95% CI]; *T. trichiura*, 0.88 [0.78 – 0.98; 95% CI]; *Taenia species*, 0.87 [0.76 – 0.98; 95% CI]; and hookworm, 0.86 [0.74 − 98; 95% CI]. Agreement among readers was good (P<.001). Among discordant results between the two readers, 6 were for *A. lumbricoides*, 5 for *T. trichiura*, 5 for *Taenia*, and 5 for hookworm. In these cases, a third reader was used to confirm the analysis.

Descriptive statistics including frequency, percentage, mean, median, range, interquartile range, and standard deviation were calculated to describe relevant variables. Cross tabulation was used to calculate the proportion of categorical variables in relation to outcome variables for any intestinal parasite infection, for *T. trichiura* infection, and for *A. lumbricoides* infection. A wealth index was constructed by using principal component analysis to code the previously listed 15 household assets as 0 (absent) or 1 (present). Internal consistency of the 15 variables was determined (Cronbach alpha of 0.78 and Kaiser-Meyer-Olkin sampling adequacy of 0.8). The socioeconomic indictors (poor, middle, and rich) were categorized based on the first component explaining 28.3% of the variance in the data with an Eigen value of 4.1.

### Multilevel, mixed-effect logistic regression for modelling infection risk

We used three data hierarchies: school-level, class-level, and individual-level (child or parent). Student participants were clustered within the same class, and classes were nested within schools. We included school and class levels during analysis and assessed potential confounding and effect modifications using multivariate, multilevel regression and stratified analysis. Prior to the multivariate regression, we checked collinearity among exposure variables. We used the presence and absence of any intestinal parasite, of *T. trichiura* infection, and of *A. lumbricoides* infection as separate outcome variables and conducted the analysis using a multilevel logistic regression model. For all predictors, we applied a simple, bivariate, logistic regression without considering a random effect and a multilevel, logistic regression model with random school and class effects.

Five models were constructed for each outcome variable (any intestinal parasite or *T. trichiura*, or *A. lumbricoides* infection). Model 1 (empty) had no covariate indicating whether to consider the random-effect model. Model II contained the individual child factors. Model III contained the individual child and individual parent factors. Model IV contained household, individual child, and individual parent factors. Finally, Model V used multilevel, multivariate, logistic regression to assess individual, household, and school factors. Exposure variables with P values <.25 in the bivariate multilevel logistic regression model were introduced into the model II, model III and model IV.

Variables in model V included individual child factors (sex, age, loss of appetite in past month, nail trimming, handwashing with soap before meals, wasting, and anemia), individual parent factors (mother’s educational status), household factors (wealth, source of drinking water, container used to store water, treated water), and school factors (participation in school feeding program). Covariate variables (e.g., sex, age, nail trimming, handwashing with soap before meals, wealth, source of drinking water, using treated water, and participation in a school feeding program) with P values >.25 in the bivariate regression model were retained in the final model to control for confounding.

Variables in model V for *T. trichiura* infection included individual child factors (sex, age, loss of appetite in past month, eating uncooked vegetable, wasting, and anemia), individual parent factors (mother’s educational status), household factors (wealth, family size, and container used to store water), and school factors (participation in school feeding program). Covariate variables (e.g., sex, age, and wealth) with P values >.25 in the bivariate regression model were retained in the final model to control for confounding. Variables in model V for *A. lumbricoides* infection included individual child factors (sex, age, nail trimming, dirt in fingernail, loss of appetite in past month, handwashing with soap after using latrine, de-worming treatment in past 6 months, and anemia), individual parent factors (mother’s educational status), household factors (wealth and source of drinking water), and school factors (participation in school feeding program). Covariate variables (e.g., sex, nail trimming, and wealth) with P values >.25 in the bivariate regression model were retained in the final model to control for confounding.

### Zero-inflated negative binomial regression for modelling infection intensity

To examine potential factors associated with infection intensity, a count model was applied using the fecal egg counts for *T. trichiura* and *A. lumbricoides* infections. A Poisson model was appropriate for count data, but the assumption of equal variance and mean did not fit to our data, because the mean of *A. lumbricoides* and *T. trichiura* eggs was higher than the variance. Moreover, one-part models tend to underestimate the frequencies of zeros and to bias estimation of the covariate effect size (38). Furthermore, our data showed about 81% (691) of excess zeroes for the *A. lumbricoides* fecal egg count and 57.7% (490) for the *T. trichiura* eggs count. The alpha dispersion parameter was significant for *T. trichiura* at 0.53 (95% CI: 0.45–0.61) and for *A. lumbricoides* infection at 0.47 (95% CI: 0.05, 0.37). Therefore, the excess zeroes in these data indicate that a zero-inflated model was appropriate. The zero-inflated negative binomial (ZINB) regression model is a two-part model that models count variables with inflated zeros and an over-dispersed count outcome (39). The ZINB model also assumes that the excess zero counts come from a logit model and the counts from a negative binomial model.

### Measure of effect and model fitness

Results were calculated as crude odds ratios and adjusted odds ratios with a 95% CI. Predictors with P values <.05 in the final multilevel, multivariate regression model were reported as statistically significant. The Vuong test was used to compare the ZINB model with a standard negative binomial model and a likelihood ratio test to compare ZINB with a zero-inflated Poisson regression model. The Vuong and likelihood ratio tests with P<.05 favored the ZINB model (38). Model fitness was checked using −2 log likelihood (deviance) and Akaike information criterion. The model with the lowest deviance and Akaike information criterion was used as the final model (38, 40).

### Ethical approval and consent to participate

The institutional review board at the College of Medicine and Health Sciences of Hawassa University (IRB/005/09) and the Regional Ethical Committee of Western Norway (2016/1900/REK vest) provided ethical clearance. The Gedeo Zone Health Department and District Education Office provided a letter of permission. School directors and teachers participated in discussions. We obtained informed written (signed) and verbal (thumb print) consent from study participants’ parents or guardians and permission (assent) from children aged 12 years and older before the interviews. The participants’ privacy and confidentiality were maintained. Children diagnosed as anemic and who tested positive for intestinal parasites were referred to the nearest health institution for treatment according to the standard national guidelines (41).

## Result

### Demographic and socio-economic status of schoolchildren and their parents

The mean age of the 861 schoolchildren (483 boys and 378 girls) was 11.4 (95% CI: 11.3–11.5) years, ranging from 7 to 14 years. About 88.4% (761/861) of mothers and 48.8% (420/861) of fathers never attended school. S1 Table summarizes the demographic and socioeconomic statuses of the schoolchildren and their parents.

### Prevalence of anemia, stunting, and wasting

Anemia occurred in 29.6% (240/810 children, 95% CI: 26.6–32.9), among whom 85% (204/240) were mildly anemic and 15% (36/240) were moderately anemic. About 32% (278/861, 95% CI: 29.2–35.5) had stunting, of which 37.4% (104/278) were severely stunted. Ten percent (85/861, 95% CI: 8.0–12.1) had wasting.

### Prevalence rates of intestinal parasites

Intestinal parasites were found in 56% (479/850) of children. Of those, 57.6% (276/479) were boys, and 30% (144/479) had multiple infections. The most frequent intestinal parasites were *T. trichiura* (75.2%; 360/479), followed by *A. lumbricoides* (33.2%; 159/479), *Taenia* species (18.2%; 87/479), hookworm species (7.7%; 37/479), *Strongyloides stercoralis* (4.4%; 21/479), and *Hymenolepis nana* (0.4%; 2/479). The mean egg intensity was 284.7 epg for *A. lumbricoides* (95% CI: 119.5–449.9), 156.4 epg for *T. trichiura* (95% CI: 127.1–185.7), 134.4 epg for *Taenia* species (95% CI: 117.4–151.4), 83.7 epg for hookworm (95% CI: 68.3–99.1), and 46.0 epg for *S. stercoralis* (95% CI: 26.0–65.9. Almost all diagnosed infections were light intensity. The S2 Table of supplementary data shows the proportions of children infected with intestinal parasites in relation to individual, household, and school factors.

*Taenia* infections were higher among children with unclean fingernails (41.4%; 36/87) than those with clean fingernails (58.6%; 51/87) (Chi square=15.3; P<.001). *Taenia* infections were also higher among children residing in households with no toilet facility (93%; 81/87) than those with a facility (7%; 6/87) (Chi square=5; P<.05). Similarly, hookworm infections were higher among children aged 10–14 years (67.6%; 25/37) than those aged 7–9 (32.4%; 12/37) (Chi square=5; P<.05). A zero epg count was observed among 81.3% (691/850) of children for *A. lumbricoides* and among 57.6% (490/850) for *T. trichiura*. The S3 and S4 supplementary tables summarize the mean, median, standard deviation, and interquartile range of *T. trichiura* and *A. lumbricoides* infections for each exposure variable.

### Risk factors for any intestinal parasite infections

Intestinal parasite infection predictors were estimated using a multivariate, multi-level, mixed-effect, logistic regression analysis. The intra-cluster correlation value, calculated in the empty model with no covariate, was 1.2% at the school and class levels and 0.1% in the final model, indicating unexplained variations of intestinal parasite infections prevalence at the school and class levels.

In the bivariate, multi-level, mixed-effect, logistic regression model, the following factors had significant associations with intestinal parasite infection: loss of appetite in the past month, wasting, anemia, having a mother or guardian with no formal education, and using an open container for water storage. In the multivariate, multi-level, mixed-effect, logistic regression model analysis, the risk of intestinal parasite infection was higher among children with loss of appetite in the past month (AOR: 1.89 [95% CI: 1.16, 3.08]), wasting (AOR: 1.73 [95% CI: (1.04, 2.90]), anemia (AOR: 1.45 [95% CI: 1.04, 2.03]), a mother or guardian with no formal education (AOR: 1.08 [95% CI: 1.25, 3.47]), and open containers for water storage (AOR: 2.06 [95% CI: 1.07, 3.99]). However, no significant differences were observed between intestinal parasite infections and sex, age, nail trimming, handwashing before meals, eating undercooked vegetables, wealth, source of drinking water, using treated water at home, or participation in a school feeding program. Table 1 and 2 shows the details

**Table 1:** Multilevel, logistic, regression analysis of predictors of intestinal parasite infection among schoolchildren in the Wonago district of southern Ethiopia, 2017

| Variables |  | Intestinal parasite |  | Adjusted OR (95% CI) |  |  |  |  |
| --- | --- | --- | --- | --- | --- | --- | --- | --- |
| Individual child factors |  | Yes (%) | No (%) | Model I | Model II | Model III | Model IV | Model V |
| Sex | Boys | 276 (57.6) | 203 (42.4) | - | 1.0 | 1.0 | 1.0 | 1.0 |
|  | Girls | 203 (54.7) | 168 (45.3) | - | .96 (.72, 1.28) | .97 (.72, 1.30) | .98 (.74, 1.32) | .98 (.74, 1.32) |
| Age in years | 7-9 | 92 (58.6) | 65 (41.4) | - | 1.19 (.82, 1.74) | 1.17 (.81, 1.71) | 1.13 (.77, 1.65) | 1.13 (.77, 1.65) |
|  | 10-14 | 387 (55.8) | 306 (44.2) | - | 1.0 | 1.0 | 1.0 | 1.0 |
| Fingernails trimmed | Yes | 386 (55.5) | 309 (44.5) | - | .85 (.52, 1.39) | .85 (.58, 1.26) | .84 (.57, 1.24) | .84 (.57, 1.24) |
|  | No | 93 (60) | 62 (40) | - | 1.0 | 1.0 | 1.0 | 1.0 |
| Dirt on fingers | Yes | 121 (58.5) | 86 (41.6) | - | 1.0 | - | - | - |
|  | No | 358 (55.7) | 285 (44.3) | - | 1.03 (.66, 1.60) | - | - | - |
| Handwashing | Always | 63 (61.8) | 39 (38.2) | - | 1.30 (.76, 2.24) | 1.31 (.78, 2.18) | - | - |
| with soap after latrine | Sometimes | 264 (54.9) | 217 (45.1) | - | 1.06 (.72, 1.56) | 1.09 (.78, 1.53) | - | - |
|  | Never | 152 (56.9) | 115 (43.1) | - | 1,0 | 1,0 | - | - |
| Handwashing before meals | Yes | 466 (56.2) | 363 (43.8) | - | .68 (.27, 1.72) | .69 (.27, 1.75) | .72 (.28, 1.83) | .72 (.28, 1.83) |
|  | No | 13 (61.9) | 8 (38.1) | - | 1,0 | 1,0 | 1,0 | 1,0 |
| Eats uncooked vegetables | Yes | 125 (60) | 83 (40) | - | 1.27 (.88, 1.82) | 1.31 (.92, 1.86) | 1.35 (.95, 1.92) | 1.35 (.95, 1.92) |
|  | No | 354 (55.1) | 288 (44.9) | - | 1,0 | 1,0 | 1,0 | 1,0 |
| Loss of appetite in past month | Yes | 82 (67.8) | 39 (32.2) | - | 1.77 (1.10, 2.85)* | 1.89 (1.18, 3.04)* | 1.89 (1.16, 3.08)* | 1.89 (1.16, 3.08)* |
|  | No | 397 (54.5) | 332 (45.5) | - |  |  |  | 1.0 |
| Body mass index-for-age (wasting) | ≥ -2 Z-score | 422 (55.0) | 345 (45.0) | - | 1.0 | 1.0 | 1.0 | 1.0 |
|  | <-2 Z-score | 57 (68.7) | 26 (31.3) | - | 1.73 (1.04, 2.88)* | 1.68 (1.01, 2.79)* | 1.73 (1.04, 2.90)* | 1.73 (1.04, 2.90)* |
| Anemia | No | 306 (54) | 261 (46) | - | 1.0 | 1.0 | 1.0 | 1.0 |
|  | Yes | 150 (63) | 88 (37) | - | 1.52 (1.09, 2.12)* | 1.49 (1.07, 2.07)* | 1.45 (1.04, 2.03)* | 1.45 (1.04, 2.03)* |
| De-worming drug past six months | Yes | 109 (57.4) | 81 (42.6) | - | .99 (.67, 1.47) | - | - | - |
|  | No | 370 (56) | 290 (44) | - | 1.0 | - | - | - |

AIC: Akaike information criterion; CI: confidence interval; NS: Not significant; OR: odds ratio; \*\*P<.01, \*P<.05

**Table 2:** Multilevel, logistic, regression analysis of predictors of intestinal parasite infection among schoolchildren in the Wonago district of southern Ethiopia, 2017

| Individual parent factors |  |  |  |  |  |  |  |  |
| --- | --- | --- | --- | --- | --- | --- | --- | --- |
| Mother's education level | Never entered school | 392 (58.5) | 278 (41.5) | - | - | 2.07 (1.26, 3.41)** | 2.08 (1.25, 3.47)** | 2.08 (1.25, 3.47)** |
|  | Read and write only | 47 (58) | 34 (41.9) | - | - | 1.97 (.97, 4.02) | 1.91 (.92, 3.97) | 1.91 (0.92, 3.97) |
|  | Primary and above | 38 (40) | 57 (60) | - | - | 1.0 | 1.0 | 1.0 |
| **Household factors** |  |  |  |  |  |  |  |  |
| Wealth | Poor | 165 (57.9) | 120 (42.1) | - | - | - | .99 (.69, 1.43) | .99 (.69, 1.43) |
|  | Middle | 165 (56.3) | 128 (43.7) | - | - | - | 1.08 (.74, 1.58) | 1.08 (.74, 1.58) |
|  | Rich | 149 (54.8) | 123 (45.2) | - | - | - | 1.0 | 1.0 |
| Source of drinking water | Unprotected | 213 (58.8) | 149 (41.2) |  | - | - | .98 (.67, 1.44) | 1.02 (.69, 1.49) |
|  | Protected | 266 (54.5) | 222 (45.5) | - | - | - | 1.0 | 1.0 |
| Water storage container | Closed container | 440 (55.3) | 356 (44.7) | - | - | - | 1.0 | 1.0 |
|  | Open container | 39 (72.2) | 15 (27.8) | - | - | - | 2.06 (1.07, 3.99)* | 2.06(1.07, 3.99)* |
| Using treated water at home | Yes | 60 (56) | 47 (44) | - | - | - | 1.07 (.68, 1.68) | 1.07 (.68, 1.68) |
|  | No | 419 (56.4) | 324 (43.6) | - | - | - | 1.0 | 1.0 |
| **School factor** |  |  |  |  |  |  |  |  |
| Participates in school food program | No | 250 (58.7) | 176 (41.3) | - | - | - | - | 1.0 |
|  | Yes | 229 (54) | 195 (46) | - | - | - | - | .98 (.66, 1.46) |
| **Variation and model fitness** |  |  |  |  |  |  |  |  |
| Variance | School level |  |  | 0.04 | 0.023 | 0.002 | 0.003 | 0.003 |
|  | Class level |  |  | NS | NS | NS | NS | NS |
| Intra-cluster correlation | School |  |  | 1.2% | 1% | 0.1% | 0.1% | 0.1% |
|  | Class |  |  | 1.2% | 1% | 0.1% | 0.1% | 0.1% |
| **Model fitness** |  |  |  |  |  |  |  |  |
|  | -2log likelihood |  |  | 1160 | 1076 | 1062 | 1057 | 1056 |
|  | AIC |  |  | 1165 | 1104 | 1089 | 1091 | 1093 |

AIC: Akaike information criterion; CI: confidence interval; NS: Not significant; OR: odds ratio; \*\*P<.01, \*P<.05

### Risk factors for *T. trichiura* and *A. lumbricoides* infections

The intra-cluster correlation value calculated in Model V for *T. trichiura* was low and insignificant, indicated that the variability in this infection prevalence was not attributable to class or school factors. The S5 Table of supplementary data shows the results of these tests. Similarly, the variability in *A. lumbricoides* prevalence at the school-level was insignificant. However, the intra-cluster correlation value calculated in Model V for *A. lumbricoides* infection indicated that 3.2% of the variability in this infection prevalence was attributable to class factors. The S6 Table of supplementary data shows the results.

All significant variables in the bivariate, multilevel, mixed-effect model also were significant in the multivariate model. The risk of *T. trichiura* infection was higher among children with loss of appetite in the past month (AOR: 1.76 [95% CI: 1.15, 2.71]), wasting (AOR: 1.73 [95% CI: 1.07, 2.78]), anemia (AOR: 1.53 [95% CI: 1.11, 2.12]), a mother or guardian with no formal education (AOR: 1.94 [95% CI: 1.18, 3.19]), and participation in the school food program (AOR: 1.55 [95% CI: 1.13, 2.12]). Furthermore, there were no statistically significant differences between *T. trichiura* infection and sex, age, nail trimming, handwashing, eating uncooked vegetable, receiving de-worming treatment in the past 6 months, or wealth. The S4 Table of supplementary data shows the results.

All significant variables in the bivariate, multilevel, mixed-effect model also were significant in the multivariate model. The odds of *A. lumbricoides* infection increased by 92% among anemic children (AOR: 1.92 [95% CI: 1.29, 2.88]). The odds were lower among children who received de-worming treatment in the past 6 months [AOR: 0.57 (95% CI: 0.33, 0.98)] and who used water from a protected source [AOR: 0.46 (95% CI: 0.28, 0.77)]. There were no statistically significant differences between *A. lumbricoides* infections and age, nail trimming, handwashing, wealth, source of drinking water, and participation in a school food program. The S5 Table of supplementary data shows the results.

### Zero-inflated negative binomial regression

#### Negative binomial count model for *T. trichiura* and *A. lumbricoides* infections

As shown in Tables 3 and 4, girls had increased intensity of *T. trichiura* infection (AOR: 1.23 [95% CI: 1.04, 1.45]). The intensity of infection with *A. lumbricoides* (AOR: 1.08 [95% CI: 1.01, 1.16]) increased with increasing age, whereas using open container for water storage at home (AOR: 1.59 [95% CI: 1.14, 2.22]) increased *T. trichiura* infection intensity. Dirty fingernails (AOR: 1.47 [95% CI: 1.07, 2.03]) were associated with increased intensity of *A. lumbricoides* infection. A habit of nail trimming (AOR: .56 [95% CI: 0.39, 0.79]) and handwashing with soap after using the latrine (AOR: 0.68 [95% CI: 0.48, 0.95]) lowered the intensity of *A. lumbricoides* infection. The epg of *A. lumbricoides* was higher among children in school feeding programs (AOR: 1.97 [95% CI: 1.49, 2.61]).

**Table 3:** Zero-inflated negative binomial regression model for *T. trichiura* fecal egg count among schoolchildren in the Wonago district, southern Ethiopia, 2017 (n=850)

| Variables |  | Zero-inflated negative binomial model |  |  |  |
| --- | --- | --- | --- | --- | --- |
|  |  |  |  | Negative binomial part | Zero-inflated part |
| Individual child factors |  | Mean (SD)<br>eggs per gram>0 | Median (IQR)<br>eggs per gram>0 | Infection intensity<br>AOR (95% CI) | Infection probability<br>AOR (95% CI) |
| Sex | Boys | 138.6 (113.5) | 120 (72-168) | 1.0 | 1.0 |
|  | Girls | 181.8 (406.2) | 120 (72-168) | 1.23 (1.04, 1.45)* | 1.06 (.79, 1.43) |
| Age in years |  | 11.4 (1.9) |  | 1.01 (.97, 1.05) | 1.07 (.98, 1.15) |
| Fingernails trimmed | Yes | 145.6 (111.4) | 120 (72-168) | .82 (.61, 1.09) | .92 (.57, 1.49) |
|  | No | 205.2 (603.2) | 108 (72-144) | 1.0 | 1.0 |
| Dirt on fingernails | Yes | 185.1 (510.6) | 120 (72-168) | 1.10 (.85, 1.43) | .87 (.56, 1.35) |
|  | No | 146.6 (114.2) | 120 (72-168) | 1.0 | 1.0 |
| Habit of eating | Yes | 146.3 (116.9) | 120 (72-168) | 1.05 (.86, 1.27) | .70 (.50, .99)* |
| uncooked vegetable | No | 160.3 (315.8) | 120 (72-168) | 1.0 | 1.0 |
| Loss of appetite in past month | Yes | 143.2 (109.5) | 120 (72-168) | .95 (.75, 1.21) | .52 (.34, .81)** |
|  | No | 159.1 (297.6) | 120 (72-168) | 1.0 | 1.0 |
| Hemoglobin concentration |  | 12.6 (1.6) |  | .96 (.91, 1.01) | 1.13 (1.03, 1.25)* |
| Body mass index-for-age (wasting) | ≥ -2 Z-score | 157.8 (292.2) | 120 (72-168) | 1.0 | 1.0 |
|  | <-2 Z-score | 147.3 (104.1) | 120 (72-168) | 1.10 (.86, 1.41) | .59 (.36, .94)* |
| Mother's education | Never entered school | 160.6 (301.2) | 120 (72-168) | 1.18(.86, 1.61) | .56 (.34,.92)* |
|  | Read and write only | 146.2 (105.9) | 120 (72-168) | 1.19 (.80, 1.78) | .64 (.32, 1.26) |
|  | Primary and above | 123.6 (70.0) | 120 (72-168) | 1.0 | 1.0 |
| Wealth status | Poor | 157.8 (130.7) | 144 (72-192) | .99 (.81, 1.22) | .98 (.68, 1.41) |
|  | Middle-class | 134.0 (85.6) | 120 (72-168) | .92 (.74, 1.14) | .95 (.66, 1.38) |
|  | Rich | 178.1 (460.2) | 96 (72-168) | 1.0 | 1.0 |
| Water storage | Closed container | 150.9 (281.7) | 120 (72-168) | 1.0 | 1.0 |
|  | Open container | 226.6 (161.2) | 168 (120-336) | 1.59(1.14, 2.22)** | .82 (.44, 1.49) |
| Participates in school food program | No | 135.7 (100.7) | 120 (72-168) | 1.0 | 1.0 |
|  | Yes | 173.9 (362.1) | 120 (72-168) | 1.18 (.98, 1.42) | .56 (.41, .78)** |
| **Model fitness** |  |  |  | **Zero-inflated Poisson regression** | **Zero-inflated negative binomial regression** |
| -2 Log-likelihood |  |  |  | 43126 | 4840 |
| Likelihood ratio test |  |  |  | - | 38000 (P < 0.001) |
| Vuong test |  |  |  | - | 17.5 (P < 0.001) |
| Akaike information |  |  |  | 43186 | 4902 |
| criteria |  |  |  |  |  |

AOR: adjusted odds ratio; CI: confidence interval; EPG: Egg per gram of stool; IQR: Interquartile ranges; SD: Standard deviation; \*\*\*P<.001 \*\*P<.01, \*P<.05

**Table 4:** Zero-inflated negative binomial regression model for *A. lumbricoides* infection eggs per gram count among schoolchildren in the Wonago district, southern Ethiopia, 2017 (n=850)

| Variables |  | Zero-inflated negative binomial model |  |  |  |
| --- | --- | --- | --- | --- | --- |
|  |  |  |  | Negative binomial part | Zero-inflated part |
| Individual child factors |  | Mean (SD)<br>eggs per gram>0 | Median (IQR)<br>eggs per gram>0 | Infection intensity<br>AOR (95% CI) | Infection probability<br>AOR (95% CI) |
| Sex | Boys | 321.6 (1213.2) | 120 (96-216) | 1.03 (.81, 1.32) | .96 (.66, 1.41) |
|  | Girls | 232.4 (609.3) | 120 (72-204) | 1.0 | 1.0 |
| Age in years |  | 11.4 (1.9) |  | 1.08 (1.01, 1.16)* | .90 (.81, .99)* |
| Fingernails trimmed | Yes | 157.8 (107.5) | 120 (72-216) | .56 (.39, .79)** | .66 (.36, 1.20) |
|  | No | 839.1 (2272.6) | 120 (96-240) | 1.0 | 1.0 |
| Dirt on fingernails | Yes | 582.5 (1797.7) | 156 (108-216) | 1.47 (1.07, 2.03)* | .64 (.37, 1.08) |
|  | No | 154.9 (112.1) | 120 (72-192) | 1.0 | 1.0 |
| Handwashing after latrine use | Always | 204 (142.0) | 132 (72-360) | .94 (.61, 1.44) | .68 (.35, 1.30) |
|  | Sometimes | 347.8 (1366.2) | 120 (72-192) | .68 (.48, .95)* | 1.39 (.85, 2.27) |
|  | Never | 240.8 (628.2) | 132 (72-216) | 1.0 | 1.0 |
| De-worming drug in past 6 months | Yes | 200 (144.5) | 120 (120-264) | 1.03 (.76, 1.41) | 1.68 (1.01, 2.78)* |
|  | No | 304.1 (1113.4) | 120 (72-192) | 1.0 | 1.0 |
| Hemoglobin |  | 12.6 (1.6) |  | .99 (.91, 1.07) | 1.20 (1.06, 1.36)** |
| concentration |  |  |  |  |  |
| Mother's education | Never entered school | 226.3 (538.4) | 120 (72-216) | 1.10 (.66, 1.85) | .45 (.21, .96)* |
|  | Read and write only | 148.2 (78.8) | 120 (96-216) | .89 (.49, 1.64) | .33 (.13, .84)* |
|  | Primary and above | 1314.7 (3557.6) | 120 (96-192) | 1.0 | 1.0 |
| Wealth status | Poor | 226.2 (512.7) | 120 (72-216) | 1.17 (.87, 1.58) | 1.17 (.74, 1.86) |
|  | Middle-class | 386 (1538.8) | 120 (72-216) | 1.17 (.87, 1.58) | 1.22(.77, 1.95) |
|  | Rich | 242.4 (665.4) | 120 (72-192) | 1.0 | 1.0 |
| Participates in school food program | No | 145.1 (79.1) | 120 (72-192) | 1.0 | 1.0 |
|  | Yes | 482.4 (1547.6) | 120 (84-228) | 1.97 (1.49, 2.61)*** | 1.37 (.87, 2.15) |
| **Model fitness** |  |  |  | **Zero-inflated Poisson regression** | **Zero-inflated negative binomial regression** |
| -2 Log-likelihood |  |  |  | 23788 | 2436 |
| Likelihood ratio test |  |  |  |  | 21000 (P<.001) |
| Vuong test |  |  |  |  | 9.4 (P<.001) |
| Akaike information criterion |  |  |  | 23845 | 2494 |

AOR: adjusted odds ratio; CI: confidence interval; EPG: Egg per gram of stool; IQR: Interquartile ranges; SD: Standard deviation; \*\*\*P<.001 \*\*P<.01, \*P<.05

#### Logit model for predicting excess zeros for *T. trichiura* and *A. lumbricoides* infections

The odds of zero epg counts for *T. trichiura* (AOR: 1.13 [95% CI: 1.03, 1.25]) and *A. lumbricoides* (AOR: 1.20 [95% CI: 1.06, 1.36]) increased with increasing hemoglobin concentrations. The odds of zero epg counts for *A. lumbricoides* eggs decreased with increasing age (AOR: 0.90 [95% CI: 0.81, 0.99]), whereas the odds increased among children who had received a de-worming drug in the past 6 months (AOR: 1.68 [95% CI: 1.01, 2.78]). The odds of zero epg counts for *T. trichiura* decreased for children who reported loss of appetite in the past month (AOR: 0.52 [95% CI: 0.34, 0.81]), wasting (AOR: 0.59 [95% CI: 0.36, 0.94]), eating uncooked vegetables (AOR: 0 .70 [95% CI: .50, .99]), a mother or guardian with no formal education (AOR: 0.56 [95% CI: 0.34, 0.92]), and participation in a school feeding program (AOR: 0.56 [95% CI: 0.41, 0.78]). However, no significant difference was observed between *T. trichiura* infections and age, nail trimming, or wealth. No statistically significant differences were observed between *A. lumbricoides* infections and sex, mother’s educational, or wealth.

## Discussion

Intestinal parasite infections were found to be a public health problem among schoolchildren aged 7 to 11 years in the Gedeo zone of southern Ethiopia. Controlling for clustering effects at the school and class levels and accounting for excess zeros of fecal egg counts, we found that intestinal parasite infections were associated with increased age, girls, wasting, anemia, loss of appetite in past month, unclean fingernails, lack of nail trimming, lack of hand washing with soap after using the latrine, de-worming treatment, mothers’ education levels, water source, and using uncovered water storage container at home. Variations attributable to both class and school-level factors for intestinal parasite infection prevalence were less than 5%, indicating minor influence.

We used a large and representative sample of schoolchildren and applied a multilevel, mixed-effect model and a ZINB model to identify risk factors for prevalence and intensity of intestinal parasite infections. Unlike previous studies, we directly involved parents of children to assess potential risk factors at the household level. We examined nutritional status and measured hemoglobin concentrations using two standard techniques, Kato-Kath and formalin-ether concentration, to enhance the detection of intestinal parasites. The dependence of clustered data within the school and class levels was measured and indicated using intra-cluster correlation. Unlike previous studies (8, 11-17), we also used a ZINB model to model infection prevalence and intensity.

Because of the cross-sectional nature of this study, causality between the outcome and the exposure variable cannot be determined with certainty. Multiple stool samples from each child could have enhanced the detection rate of intestinal parasite infection (42). Unfortunately, we did not take multiple samples, but we used two different techniques to analyze a single stool sample, which could have enhanced the detection rate. Furthermore, a species-specific diagnostic technique (43) and use of sensitive techniques, like polymerase chain reaction, could have improved the detection rate of intestinal parasite infections (44). The FLOTAC method has a higher sensitivity for hookworm detection, compared with the relatively low sensitivity of Kato-Katz and formalin-ether, and may have underestimated the prevalence of hookworm infection in this study (43, 45). We were unable to model most school-level risk factors, such as sanitation and hygiene, due to homogeneity issues.

The prevalence of intestinal parasite infections that we found was higher than previously found in southern Ethiopia (12, 24) but lower than other places (17, 46). Compared with the 24.6% prevalence found in Ethiopia (16), 26.3% in the Democratic Republic of Congo, and 26.5% found in Kenya (47, 48), we found a higher prevalence of *T. trichiura* infection. Our findings also revealed higher rates of *A. lumbricoides* infection compared with rates of 10.6% to 13% in other areas of southern Ethiopia (12, 49). The rate for hookworm infection in this study aligned with the 7.4% rate found during national mapping (49). However, it is lower than the 56.8% (16), 46.9% (50), and 18% reported in other regions of Ethiopia (24). All detected infections in this study were of light intensity (32), which is comparable with other studies in Ethiopia (24, 45, 51). Variations in intestinal parasite prevalence could be due to different diagnostic techniques and ecological settings.

Using the ZINB model, we observed increased intensity of *A. lumbricoides* infections and a decline in the probability of older children remaining free from this infection. Older children may participate in activities and environments that make them more prone to infection than younger children. In contrast, a previous study has reported a lower risk of intestinal parasite infection in the older age group (52). This reduced risk could be due to immunological and behavioral factors related to hygiene (53).

Nail and hand hygiene are well known individual factors affecting intestinal parasite infection prevalence and intensity (54). We found an increased intensity of *A. lumbricoides* infection among children with unclean finger nails, as has been reported by others (12, 15). Nail trimming and handwashing with soap after using the latrine led to reduced intensity of *A. lumbricoides* infection, similar to findings in other studies (8, 9, 14, 55). Eating uncooked vegetables has been reported as a risk factor for intestinal parasite infections (52). Our ZINB model indicated that eating uncooked vegetables lowered the probability of remaining free from *T. trichiura* infections. Ingesting contaminated raw vegetables could play an important role in transmitting intestinal parasites (56). Furthermore, receiving de-worming drugs in the past 6 months significantly increased the probability of children remaining free from *A. lumbricoides*, as has been observed in rural Bangladesh (57).

Despite a well-documented link between soil-transmitted helminth infections and undernutrition (58, 59), the evidence regarding this association varies. Some studies have reported the same risk of wasting and stunting among infected and non-infected children (11, 60, 61). In agreement with other studies (62, 63), our study revealed higher rates of intestinal parasites among wasted children. These children often lose micronutrients, which can impair nutritional status and growth (58).

The observed association between intestinal parasite infection and anemia was expected, as intestinal parasites are risk factors for anemia (64-66). Reduced food intake because of inflammatory reactions induced by lesions in the intestinal mucosa and impaired iron absorption due to worm infections could partly explain this association (58, 67). Our zero-inflation model also indicated higher probability of children remaining free from *T. trichiura* and *A. lumbricoides* infections as their hemoglobin concentrations increase. The observed infection intensity in this study was light, and although light infection by *T. trichiura* and *A. lumbricoides* may not be enough to produce significant blood loss, it may aggravate the condition (68). However, de Gier et al. found low hemoglobin concentrations among children with light *T. trichiura* infections (65). This finding could be affected by unmeasured factors, such as low dietary iron intake and malaria. Furthermore, we found high rates of intestinal parasite infection among children who reported a loss of appetite in the past month.

Intestinal parasite infection prevalence increased among children whose mothers had no formal education, similar to previous studies in Ethiopia (8, 12) and rural Mexico (69). This could be due to lack of knowledge about poor home sanitation and hygiene. Using piped water has been shown to influence the prevalence of *A. lumbricoides* infection (9). In this study, low rates of *A. lumbricoides* infection were observed among children living in households using a protected water source, indicating a possibility for contamination when water is not protected from soil-transmitted helminths eggs during transport and storage (70). We also observed a high risk of parasite infection, particularly *T. trichiura* infection, among children in households using open containers for water storage. Similar findings have been observed in Kenya (47). The high percentage of unimproved water sources and the practice of open defecation, particularly in rural Ethiopia, offer support for this finding (71). We indeed observed a high percentage of unimproved toilet facilities in the study households.

Children participating in school feeding programs had high rates of *T. trichiura* infection and increased intensity of *A. lumbricoides* infection. This finding suggests unsafe or unhygienic food preparation and poor sanitary facilities at schools in the study area. School sanitation and hygiene could affect this finding, though we were unable to show a link due to similarity of this potential exposure variable. Furthermore, some schools had no access to safe water, putting those children at higher risk. Schools with feeding programs thus may be area at high risk of food insecurity and vulnerability to infection.

Although Ethiopia launched a national school-based de-worming program in 2015, soil-transmitted helminth infection remain high among schoolchildren in the rural areas. Variations attributable to both class- and school-level factors for intestinal parasites infection prevalence were low. Most individual and few household factors were found to be important predictors for intestinal parasite infection prevalence and intensity, and high rates of *T. trichiura* infection and intensity of *A. lumbricoides* among children in school feeding programs also were observed. Interventions that improve hygiene among schoolchildren can reduce the burden of intestinal parasite infection in settings such as Gedeo. Access to safe water at school and at home is a crucial part of infection reduction strategies. Periodic de-worming programs in schools must be strengthened. To that end, school teachers should work with health workers to provide health education about personal hygiene. Integrated intervention activities focusing on the individual, household, and school will reduce the burden of intestinal parasite infections.

## Supporting information

**S1 Checklist. STROBE checklist.**

**S1 Table.** Demographic and socio-economic status of schoolchildren and their parents, Wonago district of southern Ethiopia, 2017*

**S2 Table.** Distribution of intestinal parasite infection among schoolchildren in the Wonago district, Southern Ethiopia, 2017 (n=850)

**S3 Table.** The mean, median, standard deviation, and interquartile range of *T. trichiura* infection loads in egg per gram of stool among schoolchildren in the Wonago district, southern Ethiopia, 2017 (n=850)

**S4 Table.** The mean, median, standard deviation, and interquartile range of *A. lumbricoides* infection loads in egg per gram of stool among schoolchildren in the Wonago district, southern Ethiopia, 2017 (n=850)

**S5 Table.** Multilevel logistic regression analysis of predictors of *T. trichiura* infection among schoolchildren in the Wonago district of southern Ethiopia, 2017

**S6 Table.** Multilevel logistic regression analysis of predictors of *A. lumbricoides* infection among schoolchildren in the Wonago district of southern Ethiopia, 2017

## Acknowledgements

We are grateful to the schoolchildren, parents, and guardians who participated in this study. We also thank the data collectors, supervisors, Gedeo Zone Health Department, Wonago District education office, school directors, and teachers. We are also grateful to the Ethiopian Public Health Institute for providing us the Kato-Kath template. We would like to thank Dilla University Teaching and Referral Hospital for providing us with a laboratory examination room. We are deeply grateful to Hawassa University and the University of Bergen for their support.

